# H5 influenza virus mRNA-lipid nanoparticle (LNP) vaccination elicits adaptive immune responses in Holstein calves

**DOI:** 10.1101/2025.05.01.651548

**Authors:** Carine K. Souza, Jefferson J. S. Santos, Paola Boggiatto, Haley Sterle, Bailey Arruda, Mitchell V. Palmer, Alessandra Campos, Jiaojiao Liu, Naiqing Ye, Drew Weissman, Scott E. Hensley, Amy L. Baker

## Abstract

Highly pathogenic avian influenza (HPAI) clade 2.3.4.4b H5N1 is circulating widely in lactating cows in the United States. Due to the critical need for intervention strategies for this outbreak, we evaluated antibody and cellular immune responses of a clade 2.3.4.4b H5 mRNA-LNP vaccine in calves. We found that the H5 mRNA-LNP vaccine induced a robust antibody and CD8^+^ T cellular-mediated immune response and conferred protection against clade 2.3.4.4b H5N1 infection.

## Main

Highly pathogenic avian influenza (HPAI) H5 viruses threaten wild birds, poultry and represent a human pandemic risk. In March 2024, an outbreak of HPAI clade 2.3.4.4b H5N1 virus was reported for the first time in dairy cattle^1^. Since then, over 950 HPAI H5N1 (2.3.4.4b) cases in livestock herds of dairy cattle have been confirmed, and the virus has spread to 16 states in the U.S. (http://www.aphis.usda.gov/livestock-poultry-disease/avian/avian-influenza/hpai-detections/hpai-confirmed-cases-livestock). In the field, high quantities of H5N1 have been detected in raw unpasteurized milk, although detection also occurred in conjunctivital, nasal, and urine samples of infected dairy cows^1, 2^. In experimentally intramammary inoculated cows, viral RNA was primarily detected in mammary glands and milk, although RNA was detected in a small subset of nasal and ocular swabs^3^. Most human infections associated with HPAI H5N1 clade 2.3.4.4b reported in the U.S. have been relatively mild^4, 5^. However, a severe case of HPAI H5 clade 2.3.4.4b resulted in the first U.S. H5N1-related human death (https://ldh.la.gov/news/H5N1-death). Furthermore, there is an increased concern about mammalian adaptation of the HPAI H5 clade 2.3.4.4b and the risk of human-to-human transmission^6^.

It is possible that vaccines might be able to control H5 spread in livestock and prevent the emergence of novel viruses that pose a pandemic risk to humans. Bovine H5 vaccines are needed but are currently not available. We developed a nucleoside-modified mRNA-lipid nanoparticle (LNP) vaccine expressing the hemagglutinin (HA) protein of a clade 2.3.4.4b H5 virus and found that this vaccine was immunogenic in mice and ferrets^7^. Further, the vaccine reduced morbidity and mortality in vaccinated compared to unvaccinated ferrets^7^. Here, we assessed the antibody and T cell-mediated immune responses of calves vaccinated with this clade 2.3.4.4b H5 mRNA-LNP vaccine.

For these experiments, we tested a monovalent mRNA-LNP vaccine encoding HA from the clade 2.3.4.4b A/Astrakhan/3212/2020 virus. We administered the vaccine via the intramuscular route to calves in two doses of 50 and 500 ug in a prime-boost vaccine regimen. Both H5 mRNA-LNP vaccine doses elicited antibodies that bound to the HA from the homologous vaccine strain (Fig. 1a) and the cattle 2.3.4.4b strain (Fig. 1b). Antibody levels were significantly higher in animals receiving the 500 ug dose compared to the 50 ug dose, and antibodies were boosted by the second vaccine dose in all vaccinated animals. We were also able to detect antibodies at high levels using a commercial H5 ELISA (ID Screen® Influenza H5 Antibody Competition ELISA, Innovative Diagnostics) (Fig. 1c). We found that the vaccine elicited antibodies that inhibited agglutination of a representative wild bird 2.3.4.4b strain (Fig. 1d) and neutralized both the homologous vaccine strain (Fig. 1e) and the cattle 2.3.4.4b strain (Fig. 1f).

**Figure 1.**
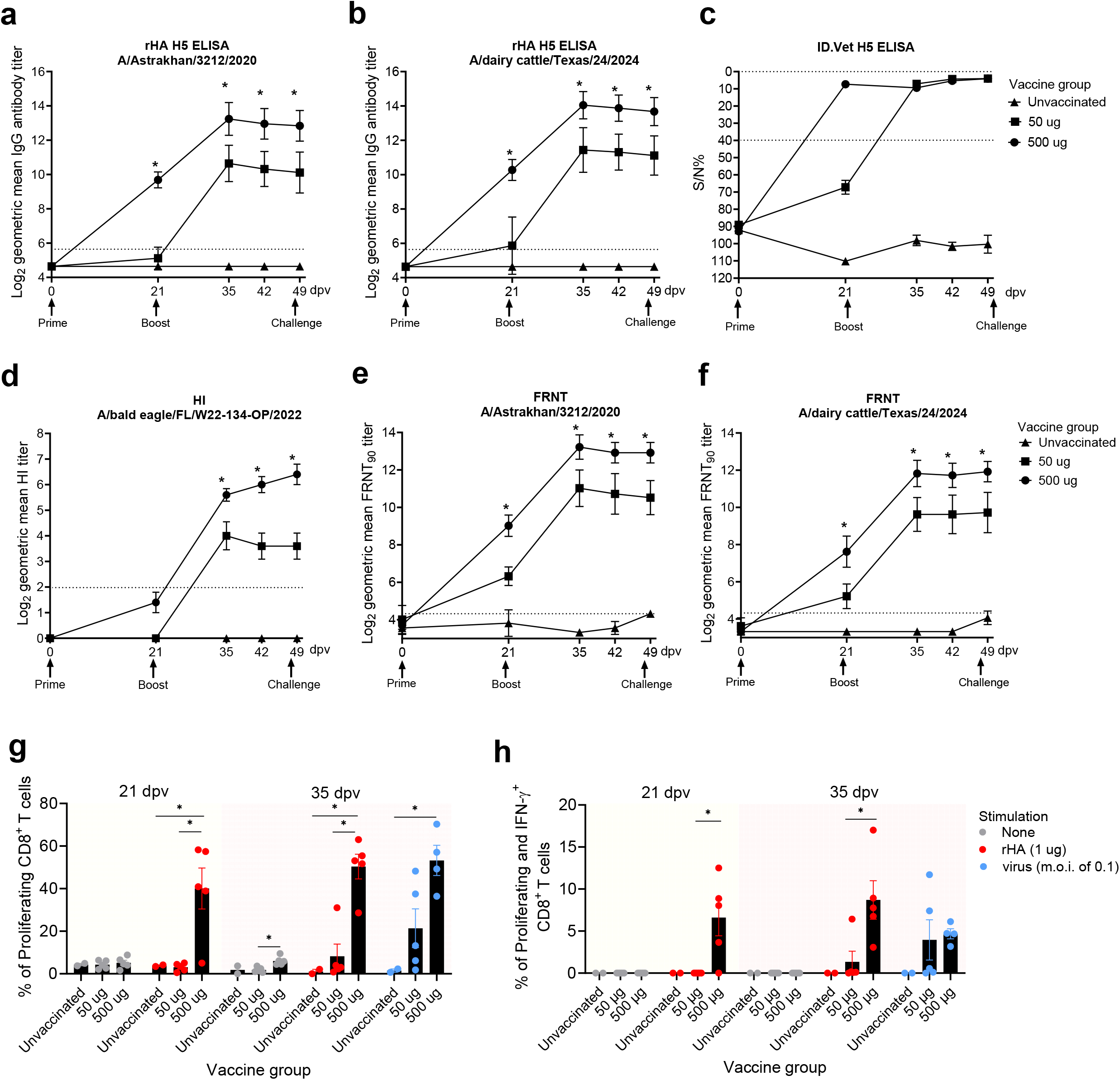
Antibody and cellular-mediated immune responses of H5 HA mRNA-LNP vaccinated calves. The *X* axis represents days post-vaccination (dpv) of the blood collected for serological assays. The *Y* axis of panels a, b, d, e and f represent the antibody titers in a Log2 scale. The Y axis of the panel c represents the SN%. rHA H5 IgG binding ELISA using the homologous vaccine virus **(a)** rHA H5 IgG binding ELISA using the 2.3.4.4b cattle strain **(b)**, commercial ID.Vet H5 ELISA for H5 antibody detection, with cut-offs of SN%<40 is positive, 40%<SN%<50% is suspect and SN%≥ 50% is negative **(c)**. Hemagglutination Inhibition assay (HI) using the wild bird representative 2.3.4.4b strain **(d)**, Foci reduction neutralization (FRNT) with the homologous vaccine strain **(e)**, and FRNT with the 2.3.4.4b cattle strain **(f)**. Percentage of proliferating CD8^+^ T cells when PBMC were stimulated with 1 ug of recombinant HA (rHA) protein homologous to the vaccine strain at 21 and 35 dpv or stimulated with m.o.i. of 0.1 of the wild bird representative 2.3.4.4b strain at 35 dpv **(g)** and the percentage of proliferating and IFN-γ^+^ CD8^+^ T cells when PBMC were stimulated with 1 ug of recombinant HA (rHA) H5 protein homologous to the vaccine strain at 21 and 35 dpv or stimulated with m.o.i. of 0.1 of the wild bird representative 2.3.4.4b strain at 35 dpv, represent the Y axis **(h)**. The unvaccinated and H5 mRNA-LNP 50 or 500 µg vaccinated groups are depicted in the *X* axis. The legend shows the PBMC non-stimulated as None in grey dots, stimulated with 1 ug of homologous recombinant HA (rHA) in red dots and stimulated with wild bird representative 2.3.4.4b strain at m.o.i. of 0.1 in blue dots. Statistical differences (p≤ 0.05) between groups and treatments are indicated by asterisks and the bar indicates group comparisons.

We next measured cell-mediated immune (CMI) responses elicited by the vaccine pre-boost at 21- and post-boost at 35-days post-vaccination (dpv) after stimulating peripheral blood mononuclear cell (PBMC) with recombinant H5 protein or virus. The 500 µg H5 mRNA-LNP vaccine dose elicited a significantly higher percentage of proliferating CD8^+^ T cells compared to the 50 µg H5 mRNA-LNP vaccine dose at 21- and 35dpv or unvaccinated, following *in vitro* stimulation with either H5 protein (Fig. 1g, red dots) or inactivated 2.3.4.4b H5 virus (Fig. 1g, blue dots). We found that CD8^+^ T cells from animals receiving the 500 ug vaccine dose produced IFN-γ^+^ after stimulation; however, CD8^+^ T cells from animals receiving the 50 ug vaccine dose only sporadically produced IFN-γ^+^ after stimulation (Fig. 1h). The vaccine also elicited low but detectable levels of proliferating CD4^+^ T cells and *γδ* T cells (Extended data Fig. 1) that produced low IFN-γ^+^ production.

We evaluated vaccine protection by feeding vaccinated and unvaccinated calves with milk from lactating cows experimentally infected with H5N1 virus via the intramammary route. Virus titers in the milk used as the inoculum ranged from 10^2.25^ - 107.5 TCID50/mL (Supplementary Table 1). Presence of viral RNA in nasal secretions of the calves were assessed via qPCR (Fig. 2a) and viral titers were detected in nasal secretions at 1-, 2-, 3-, 4- and 5-days post-infection (dpi) in the unvaccinated/challenged group. In contrast, all vaccinated calves had minimal amounts of virus detected in nasal secretions from 1 to 6 dpi (Fig. 2a). None of the calves developed a fever (Fig. 2b) or clinical signs associated with influenza virus infection. Each lung lobe homogenate was tested for presence of viral RNA by qPCR. Viral RNA was only weakly detected in one calf in the H5 mRNA-LNP 50 ug vaccine group (Supplementary Table 2). Tracheal swab and broncho-alveolar lavage fluid (BALF) samples collected at necropsy (6 dpi) contained undetectable or minimal amount of virus (Supplementary Table 2). The histopathology scores and immunohistochemistry (IHC) results from lung lobes, turbinate, and trachea samples were minimal in both vaccinated and control groups; only one calf from the unvaccinated/challenged group was positive via IHC in a single bronchiole in the caudal left lobe (Supplementary Table 3).

**Figure 2.**
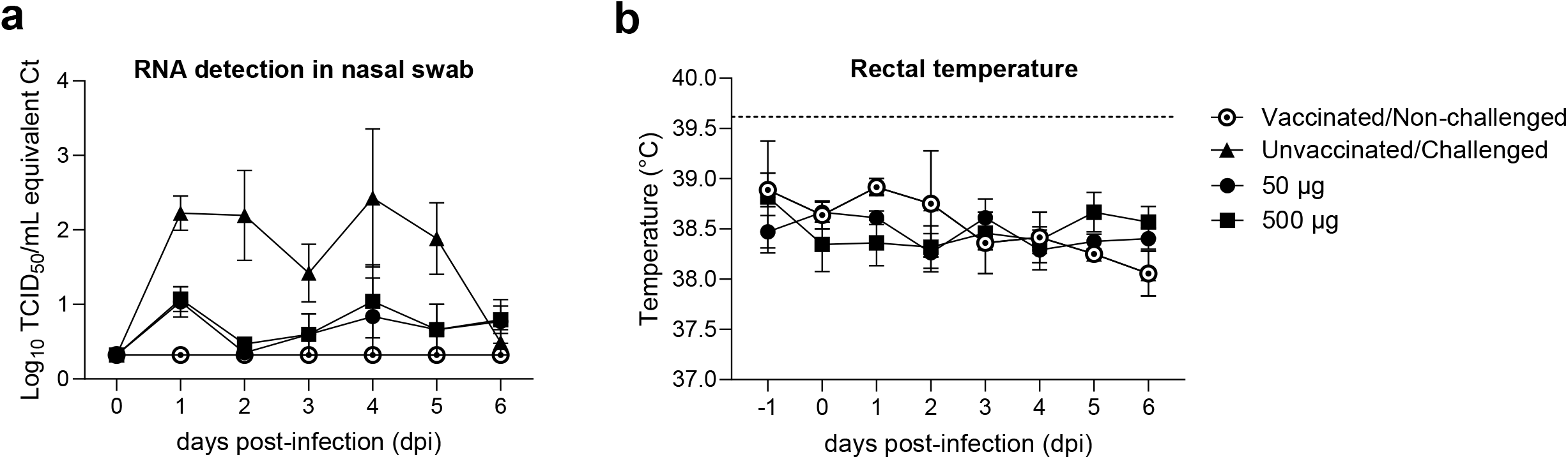
Post-challenge virus RNA detection in nasal secretions and rectal temperatures. The *Y* axis represents **(a)** the Log10 TCID50/mL equivalent Ct detection of viral RNA in nasal swabs and **(b)** rectal temperature higher than 39.7 °C indicates fever. The *X* axis represents the days post-infection (dpi).

In summary, our study demonstrates that our H5 mRNA-LNP vaccine is immunogenic in calves and reduces the amount of virus shedding following H5N1 2.3.4.4b virus infection. Future studies will evaluate this vaccine in lactating dairy cattle to determine if the vaccine provides protection against intramammary H5N1 infections. While similar levels of protection were observed for the 50 and 500 ug doses, additional experiments will be needed to determine the minimum amount of vaccine required for eliciting protective immune responses, and it will also be important to carefully determine the longevity of immune responses elicited by this vaccine. Further development of this vaccine could potentially aid in controlling the widespread H5N1 2.3.4.4b outbreak in the U.S. lactating cows.

## Methods

### Study Design

Twelve Holstein castrated male calves at 4 months of age were given 3.5 mL of Draxxin (Zoetis, Kalamazoo, MI) and 3 mL of Noromectin (Norbrook Inc., Lenexa, KS) upon arrival. At the beginning of the study, serum samples were collected and tested for Influenza A virus (IAV) antibody nucleoprotein (NP) by a commercial ELISA (MultiS ELISA, IDEXX, Westbrook, Maine) to ensure the calves were seronegative to IAV. Ten calves were split into two groups (*n*=5) and vaccinated with 50 or 500 ug of A/Astrakhan/3212/2020 (H5N8) HA mRNA-LNP vaccine intramuscularly, and two calves were kept as a non-vaccinated control group. The vaccine regimen comprised two doses of mRNA-LNP vaccine with an interval of three weeks between doses. Blood samples were collected at 0, 21, 35, 42, and 49 dpv to evaluate immune responses using serological and *in vitro* recall response assays to measure CMI responses. At 49 dpv, we challenged 4 calves from each vaccinated group and two calves as unvaccinated/challenged control group. We kept one calf from each vaccinated group as a vaccinated/non-challenged group. The challenge was performed by feeding calves, twice daily for three days, approximately 1 L of milk from cows experimentally inoculated with A/dairy cattle/Texas/24-008749-002/2024 (H5N1) virus via the intramammary route (unpublished data) ^8^. Milk samples from the inoculum were collected twice daily for three days to check virus titers by TCID50/mL virus titration was performed according to ^9^ and *Ct* values by qPCR (Supplementary Table 2). Two vaccinated/non-challenged calves were fed with non-infected milk from healthy cows.

FLOQ nylon-tipped nasal swabs (Copan, Murrieta, CA) and rectal temperatures were collected daily from 1 to 6 days post-inoculation (dpi). During the three days of challenge, nasal swabs were collected before the calves were fed with H5N1 infected milk. At 6 dpi, all calves were euthanized with a lethal intravenous dose of pentobarbital sodium (Fatal Plus, Vortech Pharmaceuticals, Dearborn, MI). Lung tissue, tracheal swab, and broncho-alveolar lavage fluid were collected at necropsy to evaluate viral RNA detection by qPCR. Lung, turbinate and tracheal tissues were also collected for histologic analysis. The animal study was carried out in an ABSL2 facility during the vaccination stage and a ABSL3-Ag facility during the challenge stage of the study in compliance with the Institutional Animal Care and Use Committee of the USDA-ARS National Animal Disease Center (NADC).

### Recombinant HA protein expression

The extracellular domain of A/Astrakhan/3212/2020(H5N8) HA and A/dairy cattle/Texas/24-008749-002-v/2024(H5N1) HA were synthetically fused to the trimeric FoldOn of T4 fibritin, the Avitag sequence, and a hexahistidine affinity tag ^10^, and cloned into the pCMV Sport6 expression vector. HA expression plasmids were co-transfected with plasmid encoding cell surfaced expressed A/Puerto Rico/8/1934 neuraminidase (NA) into FreeStyle 293F cells (Thermo Fisher, Waltham, MA) with 293fectin reagent (Thermo Fisher, Waltham, MA) following the manufacturer’s instructions. After 4 days, the supernatant was clarified by centrifugation. HA proteins were purified using Ni-NTA agarose resin (Qiagen) and gravity flow chromatography columns (Bio-Rad). Purified HA was buffer-exchanged and concentrated using centrifugal filter units with a 30 kDa molecular weight cutoff (Millipore). Purified HA proteins were stored at −80°C.

### ELISAs

ELISAs were performed on 96-well Immulon 4HBX extra high binding flat-bottom plates (Thermo Fisher, Waltham, MA). Plates were coated with HA protein at 2 μg/mL in DPBS at 4 °C overnight. Plates were washed three times with 1X PBS+0.1% Tween-20 and then blocked for 1 h at room temperature using blocking buffer (1X TBS, 0.05% Tween-20, and 1% bovine serum albumin). Bovine serum samples were heat-inactivated at 56 °C for 1 h and then serially diluted four-fold in dilution buffer (1X TBS, 0.05% Tween-20, and 0.1% bovine serum albumin). After blocking, plates were washed three times with 1X PBS+0.1% Tween-20. Diluted sera were then transferred to ELISA plates and incubated for 2 hr at room temperature (RT). After washing three times with 1X PBS+0.1% Tween-20, peroxidase-conjugated mouse anti-bovine IgG monoclonal antibody (clone IL-A2; Bio-Rad) was added to all plates at a concentration of 1:5000 in dilution buffer for 1 hr at RT. After washing three times with 1X PBS+0.1% Tween-20, all plates were developed by adding SureBlue TMB peroxidase substrate (SeraCare, Gaithersburg, MD) for 5 minutes at RT, followed by stopping the reaction with 250 mM HCl solution. The absorbance was measured at 450 nm using a SpectraMax ABS Plus plate reader (Molecular Devices, San Jose, CA). Endpoint titers were calculated as the reciprocal sera dilution that generated an equivalent optical density (OD) of 0.5.

A commercial ELISA, ID Screen Influenza H5 antibody competition 3.0 multispecies (ID Innovative Diagnostics, Grabels, France) was used to measure H5-specific antibody responses in bovine serum.

### Foci Reduction Neutralization assay

Foci reduction neutralization (FRNT) assays were carried out using conditionally replicative influenza viruses that encode GFP instead of PB1, as previously described ^11^. Conditionally replicative viruses were generated encoding the hemagglutinin (HA) and neuraminidase (NA) from either A/Astrakhan/3212/2020 (H5N8) or A/dairy cattle/Texas/24-008749-002-v/2024 (H5N1). Bovine serum samples were pretreated with receptor-destroying enzyme (RDE) (Denka Seiken, San Jose, CA) for 2 h at 37 °C, followed by heat inactivation at 56 °C for 30 min. RDE-treated sera were serially diluted in neutralization assay media (NAM, medium 199 supplemented with 0.01% heat-inactivated FBS, 0.3% BSA, 100 U of penicillin/mL, 100 µg of streptomycin/mL, 100 µg of calcium chloride/mL, and 25 mM HEPES), mixed with virus, and incubated at 37 °C for 1 h. After incubation, 2.5×10^4^ MDCK-SIAT1-CMV-PB1-TMPRSS2 cells were added to each well. Plates were incubated for 40 hours at 37 °C followed by measurement of GFP fluorescence intensity on an EnVision multi-mode microplate reader (PerkinElmer, Waltham, MA) using an excitation wavelength of 485 nm and an emission wavelength of 515 nm. Neutralization titers were expressed as the reciprocal of the highest dilution of sera that inhibited 90% of GFP expression relative to control wells with no sera.

### Hemagglutination inhibition assay

The representative wild bird 2.3.4.4b strain used in the HI assay, A/bald eagle/Florida/W22-134-OP/2022 (H5N1) is an engineered influenza viruses carrying a modified HA with a monobasic cleavage site. It is approved for work under BSL-2 conditions and was kindly provided by Richard Webby from St. Jude Children’s Research Hospital. The virus was propagated in Madin-Darby canine kidney London cell line using OptiMEM (Gibco, Thermo Scientific, Waltham, MA) supplemented with antibiotic-antimycotic (Thermo Scientific, Waltham, MA) and 1 µg/mL L-1-Tosylamide-2-phenylethyl chloromethyl ketone (TPCK, Worthington Biochemicals, Lakewood, NJ). Prior to HI, bovine 50 µL of serum was treated with 150 µL of RDE (Denka Seiken, San Jose, CA) overnight at 37 ^°^C in a water bath followed by 600 uL of saline solution (0.85%) incubated at 56 ^°^C for 30 minutes to deactivate RDE. Serum samples were adsorbed with 100% rooster red blood cells for 1 hour at 4 ^°^C to remove nonspecific hemagglutinin inhibitors and agglutinins. RDE-treated serum samples were run in HA and HI assays using 0.5% rooster red blood, slightly modified from a previously described protocol ^12^. The reciprocal titers were divided by 10, log2 transformed, and reported as the geometric mean.

### Peripheral blood mononuclear cell (PBMC) isolation and stimulation

Whole blood was collected via venipuncture of the jugular vein and placed into tubes containing ethylenediaminetetraacetic acid (EDTA) (Greiner Bio-One, Monroe, NC) to prevent clotting. PBMC were isolated via density centrifugation using Ficoll as described previously ^13^. PMBC count and viability were determined using a hemocytometer and trypan blue exclusion, and resuspended to a concentration of 1×10^7^ per ml. To assess proliferation and intracellular cytokine production, PMBC were stained with a 1:10 CellTrace Violet (eBioscience, San Diego, CA) solution, according to the manufacturer’s recommendations, and resuspended in complete RPMI media. PMBC were plated, 1×10^6^ per well, onto 96-well flat bottom plates and left unstimulated, stimulated with a purified HA recombinant protein of the homologous vaccine strain (A/Astrakhan/3212/2020) at various concentrations (0.5 ug, 1 ug, and 5 ug per well), sucrose purified, 3 x UV-inactivated wild bird representative 2.3.4.4b strain (A/bald eagle/ Florida/W22-134-OP/2022) at three multiplicity of infections (MOI) (0.01, 0.1, and 1), or concanavalin A (ConA; 0.5 ug/well; Sigma). Plates were incubated for 5 days at 37 °C with 5% CO2. Approximately 16 hours prior to harvest, all wells were treated with a 1x GolgiStop solution (eBioscience, San Diego, CA), according to the manufacturer’s recommendation.

### Flow cytometry staining and analysis

Cells were prepared for flow cytometry staining as described previously ^13^. Briefly, PBMC were transferred to 96-well round bottom plates, centrifuged at 300x g at RT, and then washed in 1x Dulbecco’s phosphate-buffered saline (DPBS). PMBC were incubated in a fixable cell viability dye solution (1:2000 dilution) (Thermo Scientific, Waltham, MA) for 20 minutes at 4 °C and then washed once in 1x DPBS and once in FACS buffer (0.5% fetal bovine serum (FBS) in PBS). PMBC were then resuspended with gentle vortexing and incubated with anti-bovine *γδ* primary antibody (Washington State University) for 15 minutes at RT and then with rat anti-mouse IgG2b secondary antibody (BUV395-labeled; R12-3) (BD Biosciences, San Jose, CA) for an additional 15 minutes at RT. PBMC were washed again twice in FACS buffer and incubated with a cocktail containing antibodies against bovine CD4 (FITC-labeled; CC30) and CD8 (AlexaFluor 647-labeled; CC58) (BioRad Antibodies, Hercules, CA) for 15 minutes at RT, followed by two additional washes in FACS buffer. For intracellular staining, PMBC were fixed and permeabilized using a permeabilization kit according to the manufacturer’s recommendations (BD Biosciences, Franklin Lakes, NJ). Intracellular cytokine staining was performed using an anti-bovine IFN-*γ* antibody (PE-labeled; CC302). PBMC were washed twice in wash buffer (BD Biosciences, Franklin Lakes, NJ), once in FACS, and then resuspended in FACS buffer until analysis. Data was collected using the BD Symphony flow cytometer (BD Biosciences, Franklin Lakes, NJ) using the DIVA software and analyzed using FlowJo®.

### Viral RNA detection in clinical samples

A piece of tissue approximately 10 mm in size was homogenized in 7.5 mL of MEM (Gibco, Thermo Fisher, Waltham, MA) in M tube using gentleMACS dissociator (Miltenyi Biotec, Auburn, CA). Lung homogenates were spun at 419 x *g* and aliquoted for viral RNA extraction. Viral RNA was extracted from 50 uL of the homogenate using Mag Max Viral RNA Kit (Thermo Fisher Scientific, MA) and reverse transcription quantitative real-time PCR (RT-qPCR) targeting IAV matrix gene, VetMAX™-Gold SIV Detection Kit (Thermo Fisher Scientific, Carlsbad, CA) were performed in the milk from the bucket samples, nasal swabs during the challenge phase and lung lobe homogenates, tracheal swab and BALF from necropsy. An aliquot of 100 uL of each lung lobe homogenate was pooled per animal and submitted to Iowa State University Veterinary Diagnostic Laboratory in Ames, Iowa, to ensure our calves were free from other respiratory pathogens such as bacteria (*Pasteurella multocida, Mycoplasma bovis, Mannheimia haemolytica* and *Histophilus somni*) or viruses (Bovine parainfluenza-3 virus, Influenza D virus, Bovine coronavirus, Bovine herpesvirus 1, Bovine viral diarrhea virus, Bovine respiratory syncytial virus) by PCR. Additionally, pooled lung homogenates were submitted four routine bacterial culture. All calves were negative for other bovine respiratory pathogens (Supplementary Table 5).

### Histologic assessment of respiratory tissues

A section of turbinate, trachea and each lung lobe (cranial part right cranial lobe, caudal part right cranial lobe, cranial part left cranial lobe, caudal part left cranial lobe, middle lobe, accessory lobe, caudal right lobe and caudal left lobe) from each calf was collected and stored in 10% formalin for 48h, then transferred to 70% alcohol. Formalin-fixed tissues were processed routinely and stained with hematoxylin and eosin (H&E). Each lung section was scored as previously described ^14^. Turbinate and trachea were scored as previously described ^15^, with slight modifications (Supplementary Table 3). Additionally, a minimum of two lung sections per animal were evaluated by immunohistochemistry (IHC) ^3, 16^. Sections stained by IHC were prioritized based on the severity of the histologic lesion score. If no histologic lesions were noted, sections from the cranial part of the left cranial lobe and middle lobe were selected. In addition, any section with a histologic score greater than 3 was stained by IHC.

### Statistical Analysis

Log2 geometric mean antibody titers and CMI responses were analyzed using analysis of variance (ANOVA), with a *P* value of <0.05 considered significant (GraphPad Prism software version 6.00; San Diego, CA). Response variables shown to have a significant effect by treatment group were subjected to pairwise comparisons using the Tukey-Kramer test, by Graph Pad Prism (10.3.0). In the post-challenge, *Ct* values from qPCR of nasal secretions, lung lobes, tracheal swabs and BALF were interpolated into a Log10 TCID50/mL standard curve of the A/dairy cattle/ Texas/24-008749-002/2024 (H5N1) strain, ranging from 10^7.75^ TCID_50_/mL to 10^-0.7^ TCID_50_/mL (Supplementary Table 4) using a simple linear regression in Graph Pad Prism (10.3.0).

## Acknowledgements

We thank the USDA NADC and NVSL leadership and personnel from animal resource and facilities and engineering units; Katharine Young, Emily Love and Sarah Anderson for laboratory technical assistance; Tonia McNunn and the NCAH select agent staff for compliance assistance; Jason Huegel, Tiffany Williams, Jonathan Gardner, Jared Peterson, Emma Hay, Emma Pratt and Brian Conrad for assistance with animal studies in the BSL-2 and BSL-3 facilities. This work was supported in part by the USDA ARS (project number 5030-32000-231-000-D); the National Institute of Allergy and Infectious Diseases, National Institutes of Health, Department of Health and Human Services (contract numbers 75N93021C00015). A.C was supported by an appointment to the USDA-ARS Research Participation Program administered by the Oak Ridge Institute for Science and Education (ORISE) through an interagency agreement between the U.S. Department of Energy (DOE) and the USDA under contract no. DE-AC05-06OR23100. Mention of trade names or commercial products in this article is solely for the purpose of providing specific information and does not imply recommendation or endorsement by the US Government. USDA is an equal opportunity provider and employer.

## Figure Legends

Extended Data 1. **Percentage of proliferating CD4**^**+**^ **and γδ**^**+**^ **T cells and IFN-γ**^**+**^ **production when PBMC were stimulated with rHA H5 protein homologous to the vaccine or the wild bird representative 2.3.4.4b virus**. The unvaccinated and 50 or 500 µg H5 mRNA-LNP vaccinated groups are depicted in the *X* axis. The Y axis represents the percentage of proliferating CD4^+^ **(a)**, IFN-γ^+^ CD4^+^ **(b)**, γ**δ**^+^ **(c)** and IFN-γ^+^ γ**δ**^+^ **(d)** T cells at 21 dpv when stimulated with rHA H5 protein homologous to the vaccine (red dots) and at 35 dpv when stimulated with rHA H5 protein homologous to the vaccine (red dots) or the wild bird representative 2.3.4.4b virus (blue dots). Non-stimulated T cells (None) are represented with grey dots. Statistical differences (p≤ 0.05) between groups and treatments are indicated by asterisks and the bar indicates group comparisons.

## Supplementary files

Supplementary Table 1. Virus titers (Log_10_TCID_50_/mL) and Ct values of the milk infected with H5N1 used as inoculum to feed calves.

Supplementary Table 2. Log_10_TCID_50_/mL equivalent Ct in lung tissues, tracheal swabs and BALF samples.

Supplementary Table 3. Histologic lesion composite scores on turbinate, trachea and lung lobes. IHC positive tissues are marked with an asterisk.

Supplementary Table 4. Log_10_TCID_50_/mL equivalent Ct and interpolated Ct values of samples.

Supplementary Table 5. Bovine respiratory pathogens panel tested from pooled lung lobe homogenate per calf.

