## supplement for "H5 influenza virus mRNA-lipid nanoparticle (LNP) vaccination elicits adaptive immune responses in Holstein calves"

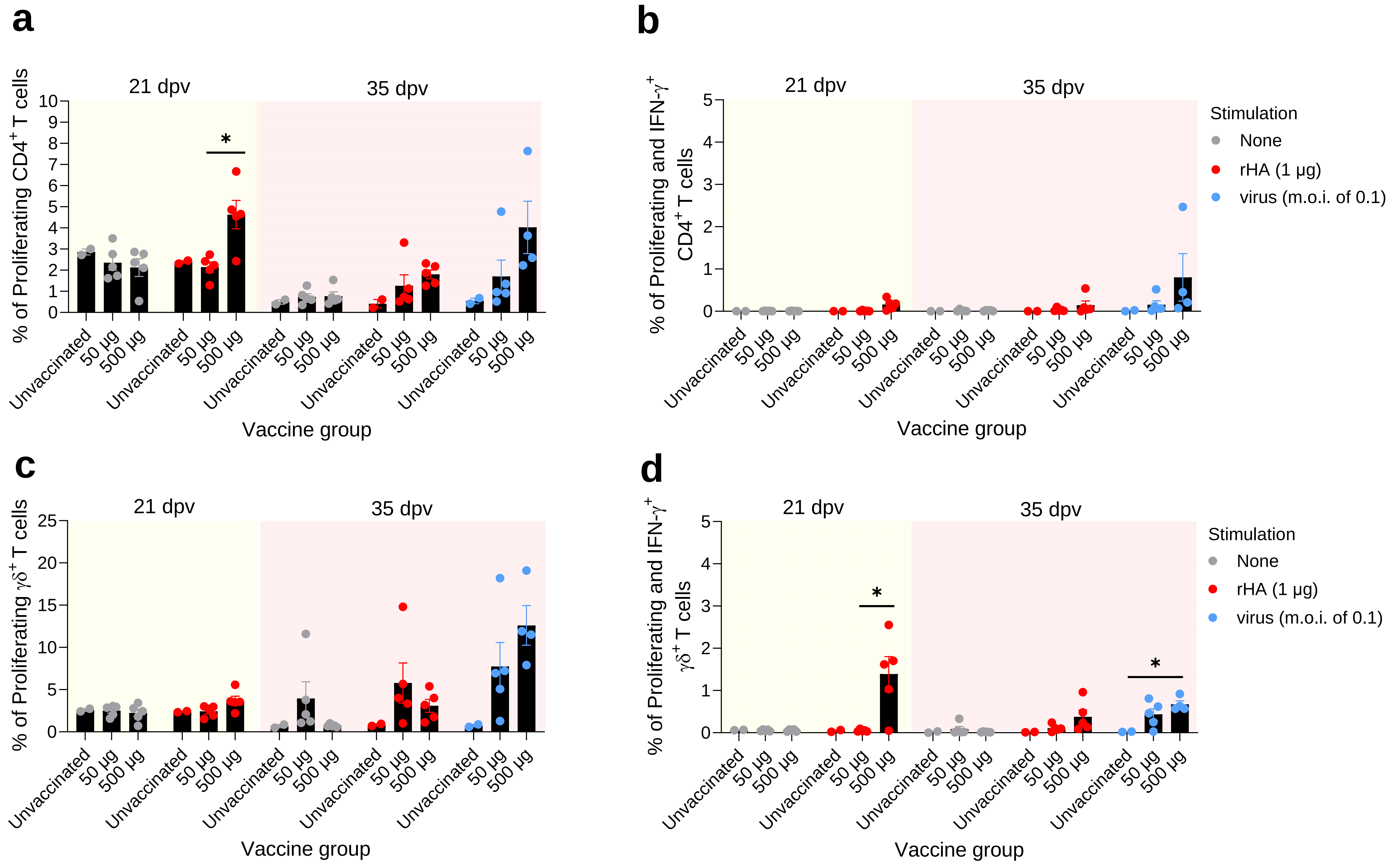

a
b
c
d
21 dpv
35 dpv
% of Proliferating CD4$^{+}$ T cells
% of Proliferating and IFN-γ$^{+}$ CD4$^{+}$ T cells
% of Proliferating γδ$^{+}$ T cells
% of Proliferating and IFN-γ$^{+}$ γδ$^{+}$ T cells
Unvaccinated
50 μg
500 μg
Vaccine group
*
Stimulation
None
rHA (1 μg)
virus (m.o.i. of 0.1)

Supplementary Table 1. Virus titers ($Log_{10}TCID_{50}$/mL) and Ct values of the milk infected with H5N1 used as inoculum to feed calves.

| DPI | Period sampled | *Ct* value | $Log_{10}$ TCID50/mL |
|---|---|---|---|
| 0 | AM | 16.4 | 7.50 |
| 0 | PM | 21.8 | 5.75 |
| 1 | AM | 26.4 | 3.50 |
| 1 | PM | 25.5 | 2.25 |
| 2 | AM | 24.7 | 2.25 |
| 2 | PM | 25.9 | 2.50 |

Supplementary Table 2. $Log_{10}$ $TCID_{50}$/mL equivalent Ct in lung tissues, tracheal swabs and BALF samples.

| | | Lung lobe | | | | | | | | | | | | | | | | | | | |
|---|---|---|---|---|---|---|---|---|---|---|---|---|---|---|---|---|---|---|---|---|---|
| | | Cranial R part cranial | | Caudal R part cranial | | Cranial L part cranial | | Caudal L part cranial | | Middle | | Acessory | | Caudal R | | Caudal L | | tracheal swab | | BALF | |
| Group | Calf# | Ct | $Log_{10}$ $TCID_{50}$/mL | Ct | $Log_{10}$ $TCID_{50}$/mL | Ct | $Log_{10}$ $TCID_{50}$/mL | Ct | $Log_{10}$ $TCID_{50}$/mL | Ct | $Log_{10}$ $TCID_{50}$/mL | Ct | $Log_{10}$ $TCID_{50}$/mL | Ct | $Log_{10}$ $TCID_{50}$/mL | Ct | $Log_{10}$ $TCID_{50}$/mL | Ct | $Log_{10}$ $TCID_{50}$/mL | Ct | $Log_{10}$ $TCID_{50}$/mL |
| | 2146 | 40.0 | 0.32 | 40.0 | 0.32 | 40.0 | 0.32 | 40.0 | 0.32 | 40.0 | 0.32 | 40.0 | 0.32 | 40.0 | 0.32 | 40.0 | 0.32 | 40.0 | 0.32 | 40.0 | 0.32 |
| Vaccinated/Non-challenged | 2154 | 40.0 | 0.32 | 40.0 | 0.32 | 40.0 | 0.32 | 40.0 | 0.32 | 39.6 | 0.45 | 40.0 | 0.32 | 40.0 | 0.32 | 40.0 | 0.32 | 40.0 | 0.32 | 40.0 | 0.32 |
| | 2143 | 40.0 | 0.32 | 40.0 | 0.32 | 40.0 | 0.32 | 40.0 | 0.32 | 40.0 | 0.32 | 40.0 | 0.32 | 40.0 | 0.32 | 40.0 | 0.32 | 40.0 | 0.32 | 40.0 | 0.32 |
| Unvaccinated/Challenged | 2151 | 40.0 | 0.32 | 40.0 | 0.32 | 40.0 | 0.32 | 40.0 | 0.32 | 40.0 | 0.32 | 40.0 | 0.32 | 40.0 | 0.32 | 40.0 | 0.32 | 40.0 | 0.32 | 40.0 | 0.32 |
| | 2144 | 40.0 | 0.32 | 40.0 | 0.32 | 40.0 | 0.32 | 40.0 | 0.32 | 40.0 | 0.32 | 40.0 | 0.32 | 40.0 | 0.32 | 40.0 | 0.32 | 40.0 | 0.32 | 40.0 | 0.32 |
| 50 ug | 2145 | 40.0 | 0.32 | 40.0 | 0.32 | 40.0 | 0.32 | 40.0 | 0.32 | 40.0 | 0.32 | 40.0 | 0.32 | 40.0 | 0.32 | 36.4 | 1.40 | 38.4 | 0.82 | 38.5 | 0.79 |
| | 2147 | 40.0 | 0.32 | 40.0 | 0.32 | 40.0 | 0.32 | 40.0 | 0.32 | 40.0 | 0.32 | 40.0 | 0.32 | 40.0 | 0.32 | 40.0 | 0.32 | 38.3 | 0.85 | 40.0 | 0.32 |
| | 2148 | 40.0 | 0.32 | 40.0 | 0.32 | 40.0 | 0.32 | 40.0 | 0.32 | 40.0 | 0.32 | 39.8 | 0.38 | 40.0 | 0.32 | 40.0 | 0.32 | 38.2 | 0.88 | 40.0 | 0.32 |
| | 2149 | 40.0 | 0.32 | 40.0 | 0.32 | 40.0 | 0.32 | 40.0 | 0.32 | 40.0 | 0.32 | 40.0 | 0.32 | 40.0 | 0.32 | 40.0 | 0.32 | 40.0 | 0.32 | 40.0 | 0.32 |
| 500 ug | 2150 | 40.0 | 0.32 | 40.0 | 0.32 | 40.0 | 0.32 | 40.0 | 0.32 | 40.0 | 0.32 | 40.0 | 0.32 | 40.0 | 0.32 | 40.0 | 0.32 | 40.0 | 0.32 | 40.0 | 0.32 |
| | 2153 | 40.0 | 0.32 | 40.0 | 0.32 | 40.0 | 0.32 | 40.0 | 0.32 | 40.0 | 0.32 | 40.0 | 0.32 | 40.0 | 0.32 | 40.0 | 0.32 | 40.0 | 0.32 | 40.0 | 0.32 |
| | 2156 | 40.0 | 0.32 | 40.0 | 0.32 | 40.0 | 0.32 | 40.0 | 0.32 | 40.0 | 0.32 | 40.0 | 0.32 | 40.0 | 0.32 | 40.0 | 0.32 | 40.0 | 0.32 | 40.0 | 0.32 |

Supplementary Table 3. Histologic lesion composite scores on turbinate, trachea and lung lobes. IHC positive tissues are marked with an asterisk.

| Group | Turbinate[a] | Trachea[b] | Lung lobe[c] | | | | | | | |
|---|---|---|---|---|---|---|---|---|---|---|
| | | | cranial part right cranial | caudal part right cranial | cranial part left cranial | caudal part left cranial | middle | acessory | caudal right | caudal left |
| Vaccinated/Non-challenged | 2.0±0.0 | 1.0±0.0 | 0.0±0.0 | 0.0±0.0 | 1.5±1.5 | 0.5±0.5 | 0.5±0.5 | 0.0±0.0 | 0.0±0.0 | 2.5±1.5 |
| Unvaccinated/Challenged | 1.5±0.5 | 1.0±0.0 | 0.5±0.5 | 1.5±0.5 | 1.0±1.0 | 1.0±1.0 | 0.5±0.5 | 2.5±1.5 | 1.5±0.5 | 2.0±2.0* |
| 50 ug | 2.3±1.0 | 1.0±0.0 | 0.8±0.8 | 0.8±0.8 | 0.5±0.5 | 0.5±0.3 | 4.0±0.6 | 1.5±0.5 | 0.0±0.0 | 1.5±0.6 |
| 500 ug | 1.5±0.9 | 0.8±0.3 | 0.3±0.3 | 0.0±0.0 | 0.3±0.3 | 0.0±0.0 | 0.0±0.0 | 0.0±0.0 | 0.0±0.0 | 0.0±0.0 |

[a] Score composite 0-8. Minimal rhinitis was defined as intraepithelial and perivascular leukocytes. Mild rhinitis: Multifocal leukocytes. Moderate rhinitis: Locally extensive leukocytes present in <3 200X HPF. Severe rhinitis: ≥ to 200X HPF and extending around submucosal glands.

[b] Score composite 0-8. Minimal tracheitis was defined as intraepithelial and perivascular leukocytes. Mild tracheitis: Multifocal leukocytes. Moderate tracheitis: Locally extensive leukocytes present in <3 200X HPF. Severe tracheitis: ≥ to 200X HPF and extending around submucosal

[c] Score composite 0-20

* IHC positive (1 calf)

Supplementary Table 4. $Log_{10}TCID_{50}$/mL equivalent Ct and interpolated Ct values of samples.

| Group | Sample | Sample type | DPI | $Log_{10}TCID_{50}$/mL | *Ct* |
|---|---|---|---|---|---|
| - | Standard 1 | - | - | 7.75 | 15.3 |
| - | Standard 2 | - | - | 6.75 | 19.1 |
| - | Standard 3 | - | - | 5.75 | 22.8 |
| - | Standard 4 | - | - | 4.75 | 26.1 |
| - | Standard 5 | - | - | 3.75 | 29.2 |
| - | Standard 6 | - | - | 2.75 | 32.6 |
| - | Standard 7 | - | - | 1.75 | 35.7 |
| - | Standard 8 | - | - | 0.75 | 38.5 |
| - | Standard 9 | - | - | 0.075 | 40 |
| Vaccinated/ Non-challenged | 2146 | nasal swab | 0 | 0.32 | 40 |
| | 2146 | nasal swab | 1 | 0.32 | 40 |
| | 2146 | nasal swab | 2 | 0.32 | 40 |
| | 2146 | nasal swab | 3 | 0.32 | 40 |
| | 2146 | nasal swab | 4 | 0.32 | 40 |
| | 2146 | nasal swab | 5 | 0.32 | 40 |
| | 2146 | nasal swab | 6 | 0.32 | 40 |
| | 2146 | tracheal swab | 6 | 0.32 | 40 |
| | 2146 | BALF | 6 | 0.32 | 40 |
| | 2154 | nasal swab | 0 | 0.32 | 40 |
| | 2154 | nasal swab | 1 | 0.32 | 40 |
| | 2154 | nasal swab | 2 | 0.32 | 40 |
| | 2154 | nasal swab | 3 | 0.32 | 40 |
| | 2154 | nasal swab | 4 | 0.32 | 40 |
| | 2154 | nasal swab | 5 | 0.32 | 40 |
| | 2154 | nasal swab | 6 | 0.32 | 40 |
| | 2154 | Tracheal swab | 6 | 0.32 | 40 |
| | 2154 | BALF | 6 | 0.32 | 40 |
| Unvaccinated/Challenged | 2143 | nasal swab | 0 | 0.32 | 40 |
| | 2143 | nasal swab | 1 | 1.99 | 34.6 |
| | 2143 | nasal swab | 2 | 1.59 | 35.9 |
| | 2143 | nasal swab | 5 | 1.03 | 37.7 |
| | 2143 | nasal swab | 4 | 3.36 | 30.2 |
| | 2143 | nasal swab | 5 | 2.37 | 33.4 |
| | 2143 | nasal swab | 6 | 0.66 | 38.9 |
| | 2143 | tracheal swab | 6 | 0.32 | 40 |
| | 2143 | BALF | 6 | 0.32 | 40 |
| | 2151 | nasal swab | 0 | 0.32 | 40 |
| | 2151 | nasal swab | 1 | 2.46 | 33.1 |
| | 2151 | nasal swab | 2 | 2.80 | 32 |
| | 2151 | nasal swab | 3 | 1.81 | 35.2 |
| | 2151 | nasal swab | 4 | 1.50 | 36.2 |

| | | | | | |
|---|---|---|---|---|---|
| | 2151 | nasal swab | 5 | 1.41 | 36.5 |
| | 2151 | nasal swab | 6 | 0.32 | 40 |
| | 2151 | tracheal swab | 6 | 0.32 | 40 |
| | 2151 | BALF | 6 | 0.32 | 40 |
| | 2144 | nasal swab | 0 | 0.32 | 40 |
| | 2144 | nasal swab | 1 | 0.82 | 38.4 |
| | 2144 | nasal swab | 2 | 0.45 | 39.6 |
| | 2144 | nasal swab | 3 | 0.32 | 40 |
| | 2144 | nasal swab | 4 | 0.32 | 40 |
| | 2144 | nasal swab | 5 | 0.32 | 40 |
| | 2144 | nasal swab | 6 | 0.32 | 40 |
| | 2144 | tracheal swab | 6 | 0.32 | 40 |
| | 2144 | BALF | 6 | 0.32 | 40 |
| | 2145 | nasal swab | 0 | 0.32 | 40 |
| | 2145 | nasal swab | 1 | 0.57 | 39.2 |
| | 2145 | nasal swab | 2 | 0.32 | 40 |
| | 2145 | nasal swab | 3 | 0.32 | 40 |
| | 2145 | nasal swab | 4 | 0.32 | 40 |
| | 2145 | nasal swab | 5 | 0.32 | 40 |
| | 2145 | nasal swab | 6 | 0.32 | 40 |
| | 2145 | tracheal swab | 6 | 0.82 | 38.4 |
| | 2145 | BALF | 6 | 0.79 | 38.5 |
| 50 μg | 2147 | nasal swab | 0 | 0.32 | 40 |
| | 2147 | nasal swab | 1 | 1.44 | 36.4 |
| | 2147 | nasal swab | 2 | 0.32 | 40 |
| | 2147 | nasal swab | 3 | 0.32 | 40 |
| | 2147 | nasal swab | 4 | 0.32 | 40 |
| | 2147 | nasal swab | 5 | 0.32 | 40 |
| | 2147 | nasal swab | 6 | 1.56 | 36 |
| | 2147 | tracheal swab | 6 | 0.85 | 38.3 |
| | 2147 | BALF | 6 | 0.32 | 40 |
| | 2148 | nasal swab | 0 | 0.32 | 40 |
| | 2148 | nasal swab | 1 | 1.31 | 36.8 |
| | 2148 | nasal swab | 2 | 0.32 | 40 |
| | 2148 | nasal swab | 3 | 1.44 | 36.4 |
| | 2148 | nasal swab | 4 | 2.40 | 33.3 |
| | 2148 | nasal swab | 5 | 1.68 | 35.6 |
| | 2148 | nasal swab | 6 | 0.88 | 38.2 |
| | 2148 | tracheal swab | 6 | 0.88 | 38.2 |
| | 2148 | BALF | 6 | 0.32 | 40 |
| | 2149 | nasal swab | 0 | 0.32 | 40 |
| | 2149 | nasal swab | 1 | 1.31 | 36.8 |
| | 2149 | nasal swab | 2 | 0.54 | 39.3 |

| | | | | | |
|---|---|---|---|---|---|
| | 2149 | nasal swab | 3 | 0.32 | 40 |
| | 2149 | nasal swab | 4 | 0.32 | 40 |
| | 2149 | nasal swab | 5 | 0.32 | 40 |
| | 2149 | nasal swab | 6 | 0.75 | 38.6 |
| | 2149 | tracheal swab | 6 | 0.32 | 40 |
| | 2149 | BALF | 6 | 0.32 | 40 |
| | 2150 | nasal swab | 0 | 0.32 | 40 |
| | 2150 | nasal swab | 1 | 0.60 | 39.1 |
| | 2150 | nasal swab | 2 | 0.32 | 40 |
| 500 µg | 2150 | nasal swab | 3 | 0.32 | 40 |
| | 2150 | nasal swab | 4 | 0.32 | 40 |
| | 2150 | nasal swab | 5 | 0.32 | 40 |
| | 2150 | nasal swab | 6 | 1.22 | 37.1 |
| | 2150 | tracheal swab | 6 | 0.32 | 40 |
| | 2150 | BALF | 6 | 0.32 | 40 |
| | 2153 | nasal swab | 0 | 0.32 | 40 |
| | 2153 | nasal swab | 1 | 1.06 | 37.6 |
| | 2153 | nasal swab | 2 | 0.69 | 38.8 |
| | 2153 | nasal swab | 3 | 0.32 | 40 |
| | 2153 | nasal swab | 4 | 1.13 | 37.4 |
| | 2153 | nasal swab | 5 | 0.32 | 40 |
| | 2153 | nasal swab | 6 | 0.32 | 40 |
| | 2153 | tracheal swab | 6 | 0.32 | 40 |
| | 2153 | BALF | 6 | 0.32 | 40 |
| | 2156 | nasal swab | 0 | 0.32 | 40 |
| | 2156 | nasal swab | 1 | 1.31 | 36.8 |
| | 2156 | nasal swab | 2 | 0.88 | 38.2 |
| | 2156 | nasal swab | 3 | 0.32 | 40 |
| | 2156 | nasal swab | 4 | 0.32 | 40 |
| | 2156 | nasal swab | 5 | 0.32 | 40 |
| | 2156 | nasal swab | 6 | 0.32 | 40 |
| | 2156 | tracheal swab | 6 | 0.32 | 40 |
| | 2156 | BALF | 6 | 0.32 | 40 |

BALF = brochoalveolar lavage fluid

Supplementary Table 5. Bovine respiratory pathogens panel tested from pooled lung lobe homogenate per calf.

| Group | Calf# | Bacterial growth | Influenza D virus | Bovine parainfluenza-3 virus | Bovine herpesvirus 1 | Bovine coronavirus | Bovine viral diarrhea virus | Bovine respiratory syncytial virus | *Pasteurella multocida* | *Mycoplasma bovis* | *Mannheimia haemolytica* | *Histophilus somni* |
|---|---|---|---|---|---|---|---|---|---|---|---|---|
| Vaccinated/Non-challenged | 2146 | Couple (Streptococcus infantarius) | Negative | Negative | Negative | Negative | Negative | Negative | Negative | Negative | Negative | Negative |
| | 2154 | Couple (no significant growth) | Negative | Negative | Negative | Negative | Negative | Negative | Negative | Negative | Negative | Negative |
| | 2143 | Few (no significant growth) | Negative | Negative | Negative | Negative | Negative | Negative | Negative | Negative | Negative | Negative |
| Unvaccinated/Challenged | 2151 | Low (no significant growth) | Negative | Negative | Negative | Negative | Negative | Negative | Negative | Negative | Negative | Negative |
| | 2144 | Negative (No aerobic growth) | Negative | Negative | Negative | Negative | Negative | Negative | Negative | Negative | Negative | Negative |
| 50 µg | 2145 | Low (Streptococcus infantarius) | Negative | Negative | Negative | Negative | Negative | Negative | Negative | Negative | Negative | Negative |
| | 2147 | Couple (Streptococcus infantarius) | Negative | Negative | Negative | Negative | Negative | Negative | Negative | Negative | Negative | Negative |
| | 2148 | Negative (No aerobic growth) | Negative | Negative | Negative | Negative | Negative | Negative | Negative | Negative | Negative | Negative |
| | 2149 | Single (No significant growth) | Negative | Negative | Negative | Negative | Negative | Negative | Negative | Negative | Negative | Negative |
| 500 µg | 2150 | Single (No significant growth) | Negative | Negative | Negative | Negative | Negative | Negative | Negative | Negative | Negative | Negative |
| | 2153 | Low (No significant growth) | Negative | Negative | Negative | Negative | Negative | Negative | Negative | Negative | Negative | Negative |
| | 2156 | Negative (No aerobic growth) | Negative | Negative | Negative | Negative | Negative | Negative | Negative | Negative | Negative | Negative |
